# Dynamical determinants of different spine movements and gait speeds in rotary and transverse gallops

**DOI:** 10.1101/2020.01.16.909481

**Authors:** Tomoya Kamimura, Shinya Aoi, Yasuo Higurashi, Naomi Wada, Kazuo Tsuchiya, Fumitoshi Matsuno

## Abstract

Quadruped gallop is categorized into two types: rotary and transverse. While the rotary gallop involves two types of flight with different spine movements, the transverse gallop involves only one type of flight. The rotary gallop can achieve faster locomotion than the transverse gallop. To clarify these mechanisms from a dynamic viewpoint, we developed a simple model and derived periodic solutions by focusing on cheetahs and horses. The solutions gave a criterion to determine the flight type: while the ground reaction force does not change the direction of the spine movement for the rotary gallop, it changes for the transverse gallop, which was verified with the help of animal data. Furthermore, the criterion provided the mechanism by which the rotary gallop achieves higher-speed than the transverse gallop based on the flight duration. These findings improve our understanding of the mechanisms underlying different gaits that animals use.

## Introduction

Quadruped animals use different gaits depending on locomotion speed, walking at low speeds and changing their gait to a trot and a canter to increase speed. In the highest range of the speed, the gait changes to a gallop. The galloping gait is generally categorized into two types: rotary and transverse gallop, involving different footfall sequences (Fig. 1) (***Hildebrand, 1977***). The rotary gallop is used by cheetahs and dogs and the transverse gallop is used by horses and bison (***Biancardi and Minetti, 2012***). Although dogs use both the rotary and the transverse gallops, the rotary gallop achieves faster locomotion than the transverse gallop (***Biancardi and Minetti, 2012***; ***Walter and Carrier, 2007***). Both gallop types involve a flight phase after the liftoff of the forelimbs (collected flight phase), in which the forelimbs and hindlimbs are collected inside while the animal is in the air (Fig. 1). However, the rotary gallop generally involves a second flight phase after the liftoff of the hindlimbs (extended flight phase), in which the forelimbs and hindlimbs are extended outside while the animal is in the air (Fig. 1a) (***Bertram and Gutmann, 2009***; ***Biancardi and Minetti, 2012***; ***Hildebrand, 1989***). While the rotary gallop has two types of flight phase, the transverse gallop has only one type of flight phase. Although there are some exceptions depending on species and conditions (e.g., dogs perform a rotary gallop with just one flight phase at low speeds (***Hudson et al., 2012***)), these characteristics of flight phases are common among species. In addition to the footfall sequence, the flight phase is also crucial for distinguishing between the rotary and transverse gallops. However, the mechanisms producing the different flight phases between the rotary and transverse galloping gaits remain unclear.

**Figure 1.**
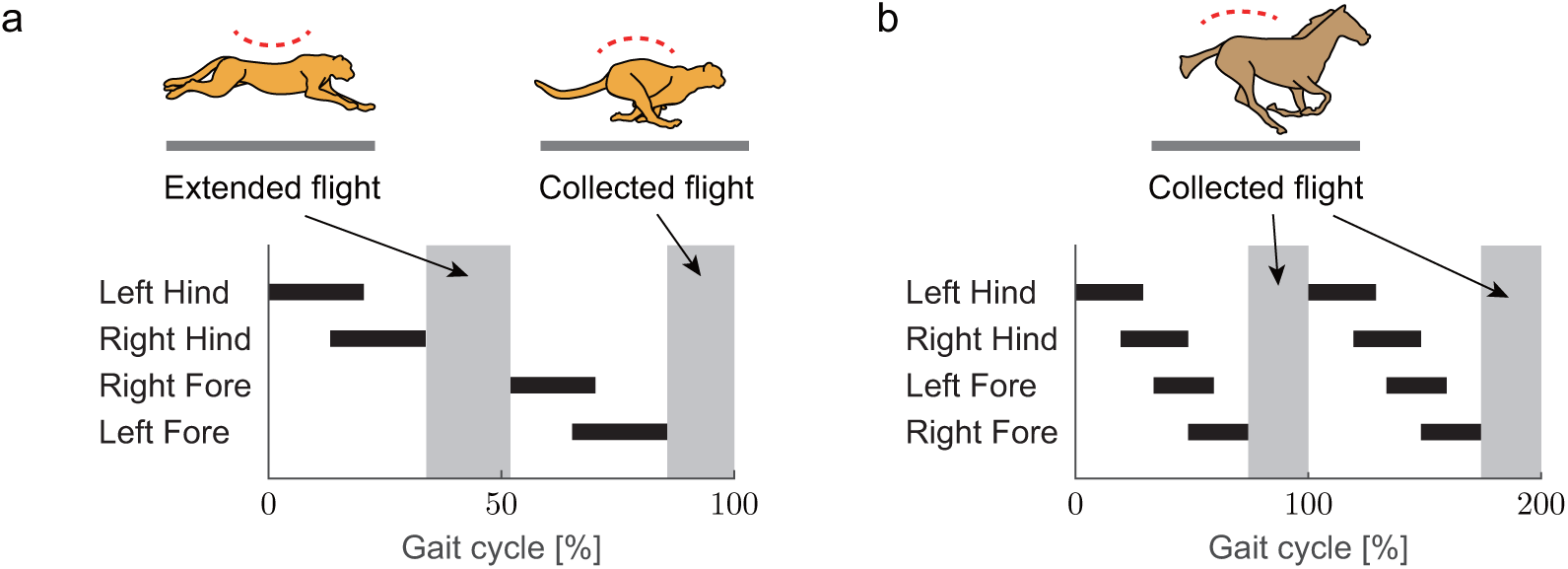
Footfall pattern and flight phases of galloping gait. (a) Cheetah rotary gallop (one gait cycle). (b) Horse transverse gallop (two gait cycles).

In galloping gait, animals flex and extend their spine during flight phases, as observed in cheetahs. Although horses exhibit only a small amount of spine bending (***Gyambaryan, 1974***), there is some motion at the sacrum (***Hildebrand, 1959***). The spine movement enhances speed by increasing the stride length, enabling the limbs to swing further (***Walter and Carrier, 2007***; ***Hildebrand, 1959***; ***Schilling and Hackert, 2006***; ***Hildebrand, 1961***; ***English, 1980***). Moreover, this movement improves energy efficiency because the energy is stored in the elastic elements of the body then released (***Taylor, 1978***; ***Alexander, 1988***; ***Minetti et al., 1999***). Spine movement differs between the two types of flight phase. Specifically, while the spine is flexed in collected flight (Fig. 1a), it is extended in extended flight (Fig. 1a). This difference of spine movement is crucial for distinguishing between the rotary and transverse gallops.

Based on observational data, ***Biancardi and Minetti*** (***2012***) suggested that animals demonstrating a preference for the transverse gallop generally have heavier bodies and longer proximal limb segments compared with animals that tend to use the rotary gallop. That is, selection between the two galloping gaits appears to depend on biomechanical determinants. However, animal running is governed by two different dynamics in the flight and stance phases. During the flight phase, all feet are in the air and the center of mass (COM) of the whole body exhibits ballistic motion. In contrast, during the stance phase, some of the feet are in contact with the ground and the legs behave like springs. Because of the complex nature of the governing dynamics, there are limitations to the understanding of the mechanisms underlying different flight phases in the galloping gait types that can be gained from observations of animals alone. To overcome the limitations of the observational approach in animal locomotion, modeling approaches have attracted recent research attention (***Bertram and Gutmann, 2009***; ***Alexander, 1988***; ***Marques et al., 2014***; ***Markowitz and Herr, 2016***; ***Swanstrom et al., 2005***; ***Aoi et al., 2017***; ***Usherwood and Davies, 2017***; ***Ambe et al., 2018***; ***Fujiki et al., 2018***; ***Aoi et al., 2019***; ***Toeda et al., 2019***). Because the essential contribution of the legs can be represented by springs, the spring-loaded inverted pendulum (SLIP) model was developed to investigate animal locomotion mechanism from a dynamic viewpoint, particularly for human running and walking (***Blickhan, 1989***; ***McMahon and Cheng, 1990***; ***Seyfarth et al., 2002***; ***Geyer et al., 2005, 2006***; ***Srinivasan and Holmes, 2008***; ***Lipfert et al., 2012***; ***Seethapathi and Srinivasan, 2019***). The SLIP model has been improved for examining gait in quadrupeds (***Full and Koditschek, 1999***; ***Blickhan and Full, 1993***; ***Farley et al., 1993***; ***Tanase et al., 2015***) to clarify the common and unique principles between animal gaits. Recently, the SLIP model has been further improved to investigate the dynamic roles of spine movement in quadruped running (***Cao and Poulakakis, 2013***; ***Pouya et al., 2017***; ***Wang et al., 2017***; ***Kamimura et al., 2015, 2018***). However, the mechanisms underlying the different flight phases between the rotary and transverse gallops remain unclear.

To clarify the mechanisms, we developed a simple analytical model based on an improved SLIP model to derive periodic solutions by focusing on the rotary gallop of cheetahs and the transverse gallop of horses. We achieved the solutions with two different flight types and spine movements corresponding to the rotary gallop, and solutions with only one flight type corresponding to the transverse gallop. We obtained a criterion to determine the solution type: while the ground reaction force (GRF) does not change the direction of the spine movement for the cheetah rotary gallop, it changes for the horse transverse gallop, which was verified with the help of animal data. The criterion also provided the mechanism by which the rotary gallop produces higher-speed loco-motion than the transverse gallop. We discussed the mechanisms for different spine movements and gait speeds in rotary and transverse gallops based on our findings.

## Materials and methods

### Model

In this study, we used a two-dimensional physical model consisting of two rigid bodies and two massless bars (Fig. 2). The bodies are connected by a joint, which is modeled to emulate the spine bending movement, and has a torsional spring with a spring constant of *K*. The bars represent the legs. We assumed that the fore and hind parts of the model have the same physical parameters. *X* and *Y* are the horizontal and vertical positions, respectively, of the COM of the whole body. 2*ϕ* is the spine joint angle. The mass and moment of inertia around the COM of each body are *M* and *J*, respectively. The lengths of each body and leg bar are 2*L* and *H*, respectively. The distance between the COM and root of the leg bar is *D*, which is positive when the root is outside the COM relative to the spine joint. The torsional spring is at the equilibrium position when the fore and hind bodies are in a straight line (*ϕ* = 0). *g* is the gravitational acceleration. In this study, we used the same model for cheetahs and horses but used different values for the physical parameters.

**Figure 2.**
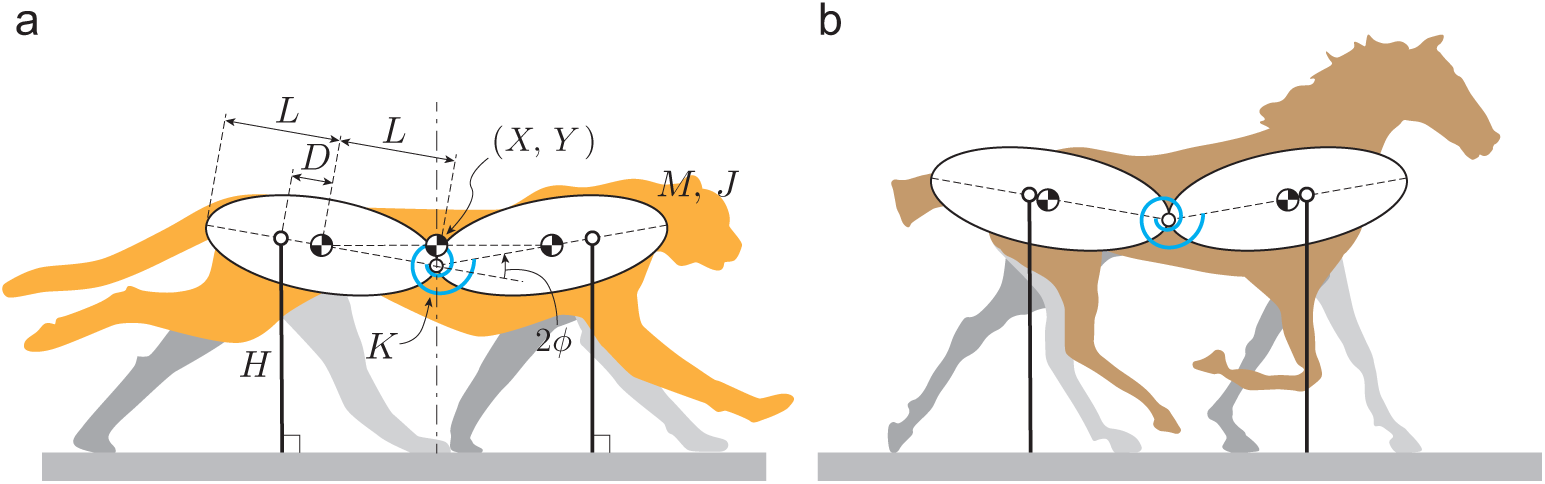
Model for (a) cheetah and (b) horse composed of two rigid bodies and two massless bars. The bodies are connected by a joint with a torsional spring. The bars represent the legs. The physical parameters are different between the models of the cheetah and horse.

In animal galloping, the pitching movement of the line connecting the root of the neck and the hip is relatively small. In our previous study (***Kamimura et al., 2015***), we used a physical model composed of two rigid bodies and two legs, which was able to perform a pitching movement as well as vertical movement. The simulation results revealed that the vertical movement of the COM and the spine joint angle between the bodies were significant determinants of the dynamic characteristics of bounding gait, compared with pitching movement. In the current study, we neglected the pitching movement of the model, making the COM vertical positions of the fore and hind bodies identical. Furthermore, even when we also ignored the horizontal ground reaction force in our previous work (***Kamimura et al., 2018***), the principal dynamic characteristics in bounding gait remained unchanged. Therefore, we also neglected the horizontal ground reaction force of the model, which allowed us to ignore the dynamics of *X*, and assumed that the leg bars were always vertical to the ground. We discuss the effects of our assumption about the pitching movement on the galloping dynamics in more depth in Supplementary information S1.

During the flight phase, the equations of motion for *Y* and *ϕ* are given by

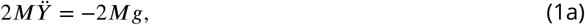

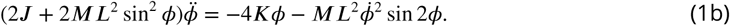

When the foot touches the ground, it receives the GRF. Because the COM vertical positions are identical between the fore and hind bodies, the foot contact of the fore and hind legs occurs simultaneously. This condition is given by

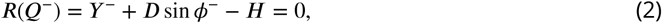

where 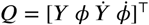 and *^−^ indicates the state immediately prior to the foot contact. Because the duty factor in animal galloping is small (***Hudson et al., 2012***), we assumed that the stance phase is sufficiently short and that the foot contact can be regarded as an elastic collision, involving no position change and energy conservation. The relationship between the states immediately prior to and immediately following the foot contact is given by

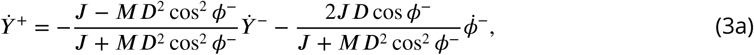

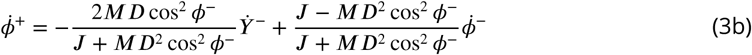

where *^+^ indicates the state immediately following foot contact. The derivation of these equations is presented in Supplementary information S2.

In this study, we solved these governing equations under the condition |*ϕ*| ≪ 1 and 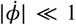. The linearization of the equations of motion (1) gives

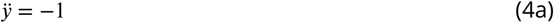

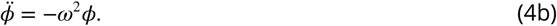

where 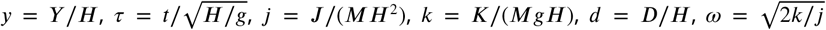, and from now on, 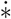 indicates the derivative of variable ∗ with respect to *τ*. The foot-contact condition (2) is approximated by

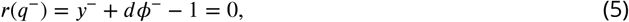

where 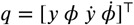. The foot-contact relationship (3) is linearized by

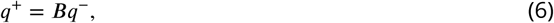

where

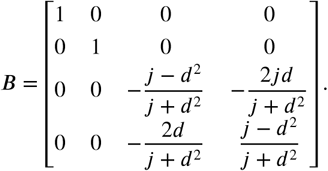

### Derivation of periodic solution

Rotary galloping involves two flight phases and two stance phases in one gait cycle. In this study, we obtained analytical periodic solutions with two flight phases and two foot contacts for each gait cycle based on the linearized equations (4), (5), and (6) (because transverse galloping has two flight phases and two stance phases in two gait cycles, we assumed this as one gait cycle for the solutions).

We defined the periodic solution as 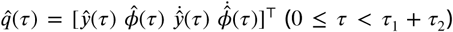, where *τ* = 0 is the onset time of the first flight phase and *τ*_1_ and *τ*_2_ are the durations for the first and second flight phases, respectively. From (4), we obtain

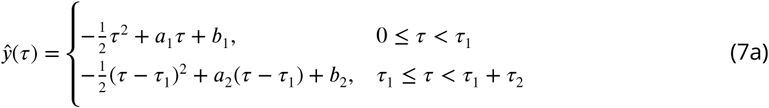

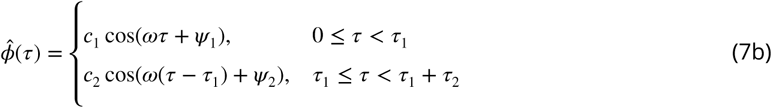

where *a*_*i*_, *b*_*i*_, *c*_*i*_ > 0, and −*π* ≤ *Ψ*_*i*_ < *π* (*i* = 1, 2) are constant. We assumed that *τ*_*i*_ < 2*π*/*ω* (*i* = 1, 2) because animals do not oscillate their spines more than once in one gait cycle. To obtain the periodic solution, we have to determine *a*_*i*_, *b*_*i*_, *c*_*i*_, *Ψ*_*i*_, and *τ*_*i*_ (*i* = 1, 2). *c*_1_ and *c*_2_ indicate the amplitudes of the first and second spine joint oscillations, respectively.

Because the foot-contact condition is satisfied at the first foot contact (*τ* = *τ*_1_) and second foot contact (*τ* = *τ*_1_ + *τ*_2_), (5) gives

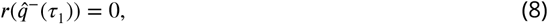

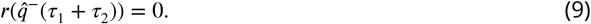

From the foot-contact relationship (6) and periodic condition, we obtain

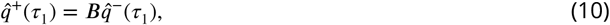

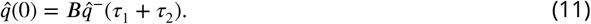

From the conditions (8)-(11), we determine ten constants *a*_*i*_, *b*_*i*_, *c*_*i*_, *Ψ*_*i*_, and *τ*_*i*_ (*i* = 1, 2) to obtain the periodic solution. However, these conditions produced various types of solutions, including solutions that are unlikely in animals. Therefore, we focused on solutions which satisfy

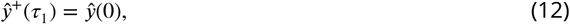

so that the COM vertical position remained unchanged at each foot contact, as shown in Fig. 3a. Under this assumption, the periodic solution is symmetric with respect to *τ* = *τ*_1_/2 and *τ* = *τ*_1_ + *τ*_2_/2 from the periodic condition, as shown in Fig. 3b. It has been reported that quadruped animals show this symmetric property in locomotion (***Raibert, 1986***). The symmetry condition (12) forces the third and fourth rows in (11) to be satisfied and reduces two conditions (this mechanism is presented in Supplementary information S3). As a result, the number of independent conditions is reduced to nine. To find a unique solution, another condition (e.g., total energy) is needed.

**Figure 3.**
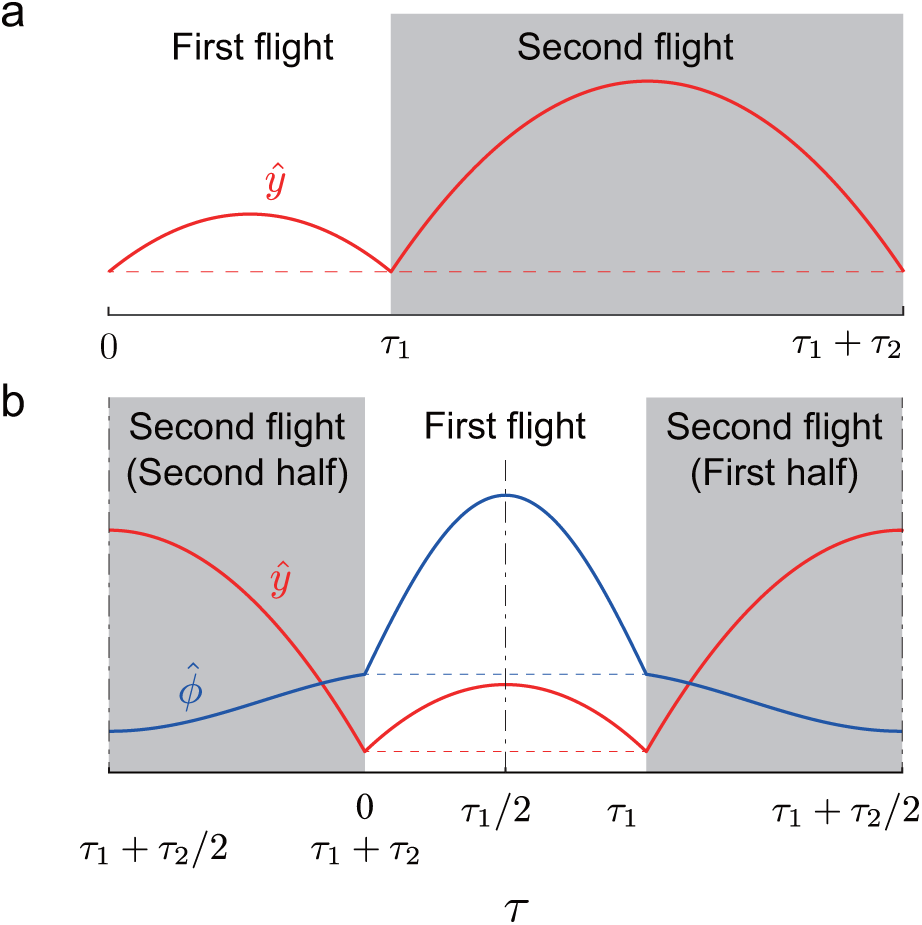
Symmetry condition for periodic solution. (a) *ŷ*^+^(*τ*_1_) = *ŷ*(0) for *ŷ*(*τ*). (b) Symmetric periodic solutions with respect to *τ* = *τ*_1_/2 and *τ* = *τ*_1_ + *τ*_2_/2

### Classification of solutions

The flight phases are classified into two types based on the spine joint movement: collected and extended. In collected flight, the spine joint is flexed (*ϕ* < 0) at the mid-flight phase. Therefore, 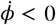 is satisfied at the beginning of collected flight. In extended flight, the spine joint is extended (*ϕ* > 0 at the mid-flight phase. Therefore, 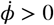 is satisfied at the beginning of extended flight. Because periodic solutions have two flight phases in one gait cycle, periodic solutions are classified into four types (both collected, both extended, first collected and second extended, and first extended and second collected). In addition, some periodic solutions have two identical flights, which satisfies 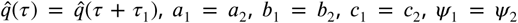, and *τ*_1_ = *τ*_2_. We distinguished such solutions from those which had two different flights, and classified the solutions into two types (same two collected and same two extended). The solutions of these two types are identical to those obtained in our previous work (***Kamimura et al., 2018***).

As a result, the periodic solutions are classified into the following six types, as shown in Fig. 4:

**Figure 4.**
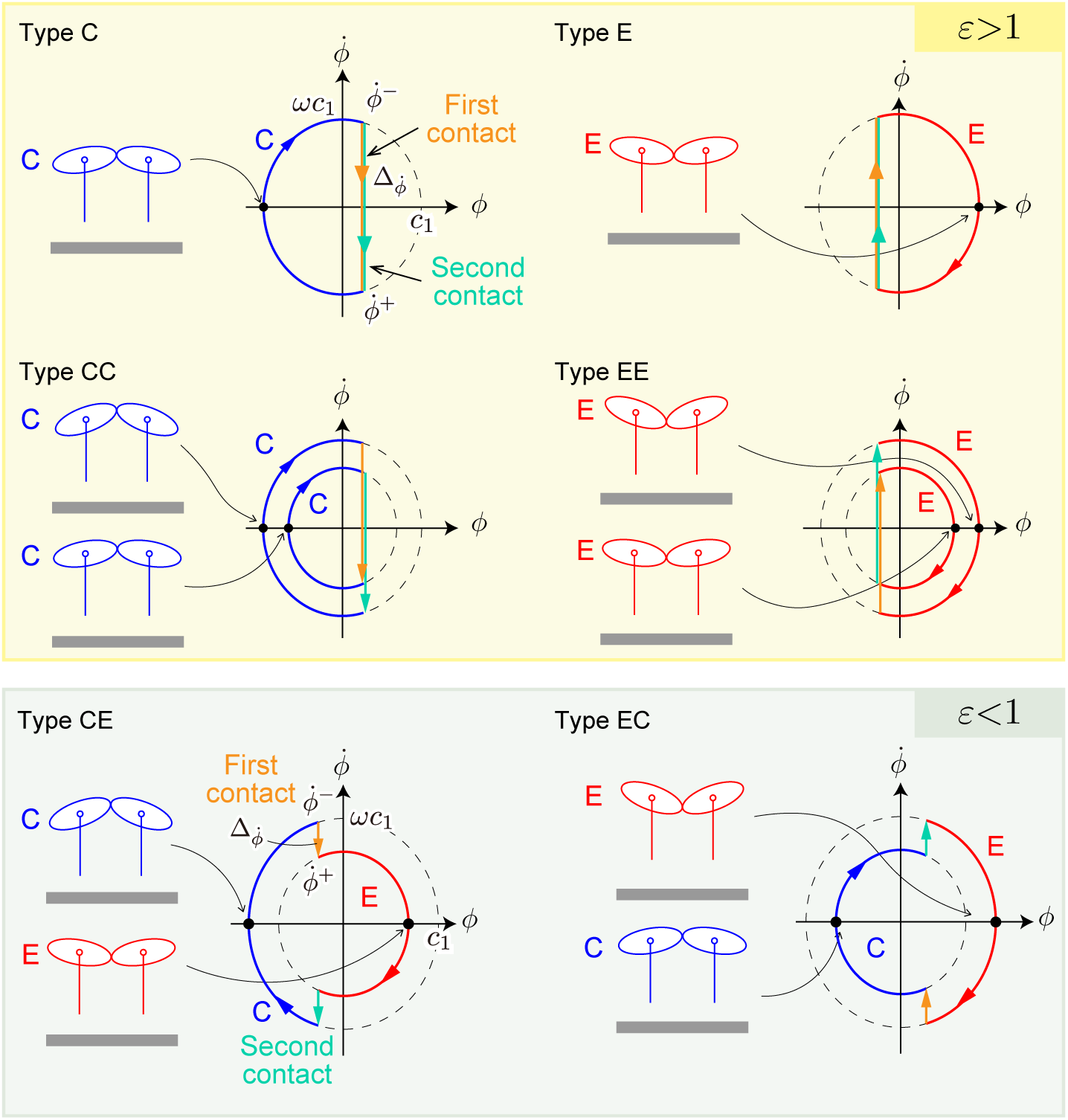
Six types of solutions. C and E indicate collected and extended flights, respectively. 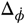 is the difference of 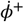 and 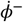, which indicate angular velocities immediately prior to and following foot contacts, respectively. *ωc*_1_ is amplitude of angular velocity. *ε* is ratio of the angular velocity change 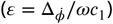. When *ε* > 1, solutions have one type of flight (solutions of type C, E, CC, and EE). Solutions of type CE and EC satisfy *ε* < 1.

1. Type C: Same two collected flights
2. Type E: Same two extended flights
3. Type CC: Two different collected flights
4. Type EE: Two different extended flights
5. Type CE: Two different flights (first: collected, second: extended)
6. Type EC: Two different flights (first: extended, second: collected)

We distinguished types CE and EC with the assumption that the amplitude of oscillation of *ϕ* in the first flight phase is greater than that of the second flight phase (*c*_1_ > *c*_2_).

The periodic solution gives an important criterion to determine the solution type; the signs of 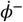 and 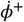 are different for one type of flight (solutions of types C, E, CC, and EE) while they are identical for two different types of flights (solutions of types CE and EC). The criterion gives a hypothesis: while the effect of GRF is too small to change the direction of the spine movement in the gallop with two different types of flights, the effect is so large that the direction changes in the gallop with one type of flight. We evaluated this hypothesis as follows. The difference of 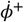 and 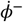 is given by

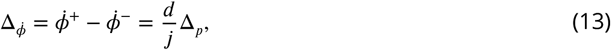

where 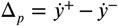 is the vertical impulse at the foot contact. To investigate the ratio of the angular velocity change to the amplitude of the angular velocity, we define

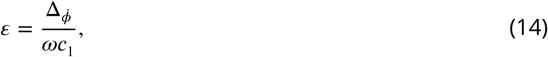

When *ϵ* > 1, solutions have one type of flight. In contrast, solutions with two different flight types have *ϵ* < 1.

Small gait cycle durations allow animals to kick the ground frequently for acceleration and achieve high-speed locomotion (***Hudson et al., 2012***). Because short flight durations induce small changes in the COM height, we investigated the COM height changes. From (7), we obtained the COM height changes in the first flight *δh*_1_ and second flight *δh*_2_ by the difference between the maximum height at apex and the minimum height at foot contact in each flight as follows:

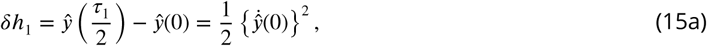

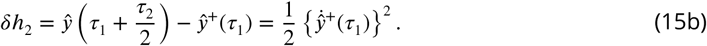

### Stability analysis

When we found periodic solutions, we computationally investigated the local stability from the eigenvalues of the linearized Poincaré map around the fixed points on a Poincaré section. We defined the Poincaré section by the state just after the second foot contact. Because our model is energy conservative, the gait is asymptotically stable when all the eigenvalues except for one eigenvalue of 1 are inside the unit cycle (these magnitudes are less than 1).

### Measurement of animals

To determine the physical parameters (*M, L*, and *J*) of our model for cheetahs, we used whole-body computed tomography (CT) from one fresh cadaver of an adult cheetah, obtained from Himeji Central Park (Hyogo, Japan). A total of 1,941 consecutive cross-sectional images were obtained using a Supria scanner (Hitachi Medical Corporation, Tokyo, Japan) at the Yamaguchi University Animal Medical Center. The tube voltage and current were set to 120 kV and 200 mA, respectively. The pixel size of each image was 0.841 mm and the slice interval was 0.625 mm. Observations of the spinal oscillation of cheetahs during galloping indicate that the anti-node is located around at the twelfth thoracic vertebra (T12). Therefore, we divided the CT into the fore and hind parts at T12. We calculated the physical parameters at each part individually and averaged them. To calculate the mass *M*, COM position *L*, and moment inertia *J* around the COM, we approximated the body as multiple elliptical cylinders and assumed that the density is uniform and 1,000 kg/m^3^. To determine these parameters for horses, we used measured data from ***Swanstrom et al***. (***2005***) and ***Grossi and Canals*** (***2010***).

The length of the leg bar *H* indicates the height of the leg root during the stance phase of galloping, and is different from the actual leg length. To determine *H*, we used kinematics data of cheetahs and horses measured during galloping. We used four adult male cheetahs (40–50 kg) at Shanghai Wild Animal Park (Shanghai, China), who were raised from infancy at the zoo. All experimental protocols were approved by the Institutional Animal Care and Use Committee at Yamaguchi University. The cheetahs were encouraged to run around a 400 m track at the zoo using a lure that traveled ahead of them at a speed of 15 to 18 m/s. Their motion was measured from the lateral side using six high-speed cameras (EX-F1 cameras, CASIO, Tokyo, Japan) at a sampling rate of 600 Hz. We used eight strides during straight running (five from one cheetah and three from the others). We used one adult male thoroughbred horse (approximately 500 kg, 6 years old) at Japan Racing Association (Tokyo, Japan). The horse was encouraged to run on a motorized treadmill (Mustang 2200, Kagra AG, Fahrwangen, Switzerland). The motion was measured from the lateral side using one camera. We used four strides during running on the treadmill. We also used the photographs of one stride of five horses galloping in ***Muybridge*** (***1957***). We determined *H* from the average of the heights of the shoulder joint and the greater trochanter of the femur during the stance phase for both cheetahs and horses.

Our model received an impulsive force at the foot contact and the distance *D* determines the position in the model to receive the force. To determine *D*, we used the vertical GRF data of cheetahs and horses measured during galloping in ***Hudson et al***. (***2012***) and ***Niki et al***. (***1984***), respectively, as well as the kinematics data above. Specifically, we first calculated the percentage of the stance phase when half of the net impulse during the stance phase was applied in each fore leg and hind leg. We then calculated the horizontal positions of the toe and leg root at the moment from the measured kinematics data in the fore legs and hind legs individually and averaged them to determine *D*.

We determined the spring constant *K* from 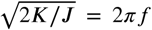, where *f* is the stride frequency determined from the value estimated in ***Hudson et al***. (***2012***) for cheetahs and the value estimated in ***Minetti et al***. (***1999***) for horses.

We used the measured amplitudes of the spine oscillation to evaluate the obtained solutions. Specifically, we compared the first amplitude *c*_1_ and second amplitude *c*_2_ of stable solutions of the types CE and EC in the cheetah model with the measured amplitudes during the first and second flights of cheetahs. In contrast, we compared the amplitude *c*_1_ of stable solutions for types C and E (*c*_1_ = *c*_2_) in the horse model with measured amplitudes during the flight phases of horses.

To estimate the ratio of the angular velocity change *ϵ* in (14), we calculated the impulse Δ_*p*_ from the measured GRF data in ***Hudson et al***. (***2012***) and ***Niki et al***. (***1984***) for cheetahs and horses, respectively. Specifically, we calculated the vertical impulses of each leg individually and averaged them as 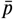. While we determined 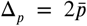 for cheetahs because two legs touch the ground in one stance phase, and 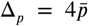 for horses because four legs touch the ground in one stance phase (Fig. 1). We determined *c*_1_ from the measured amplitude of the first flight and *ω* from 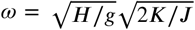.

We estimated the COM height changes from the measured kinematic data of cheetahs and horses; COM height changes were obtained by the difference between the maximum height at apex in each flight and the averaged minimum height of the mid-stances of fore legs and hind legs. For cheetahs, the collected flight was used for the first flight h_1_ because the spine is mainly bent during collected flight.

## Results

### Periodic solutions and comparison with measured data

To determine the periodic solution in (7), we obtained *a*_1_, *a*_2_, *b*_1_, *b*_2_, *c*_2_, *Ψ*_2_, *τ*_1_, and *τ*_2_ from (8)-(12) as functions of *Ψ*_1_ and *c*_1_ as follows:

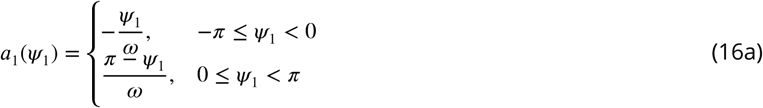

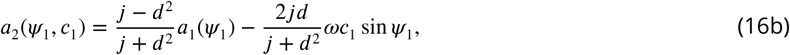

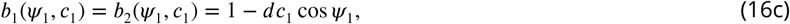

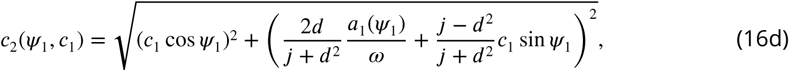

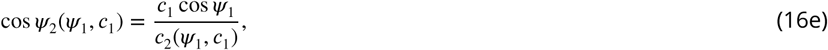

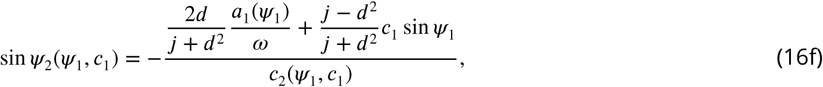

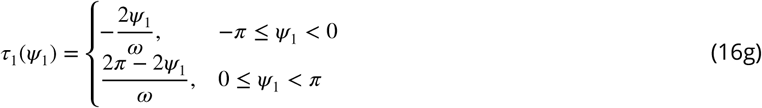

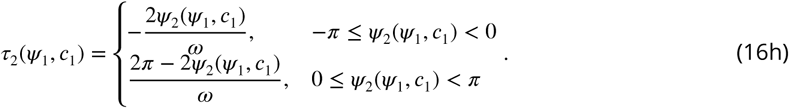

*ψ*_1_ and *c*_1_ satisfy Γ (*ψ*_1_,*c*_1_) = 0 where

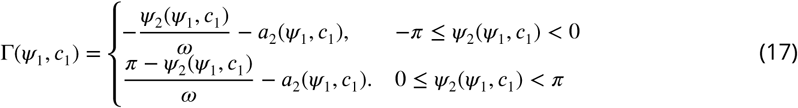

These functions depend on *j, d*, and *k* (this appears as 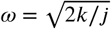).

From the measured data of cheetahs, we determined *M* = 19 kg, *J* = 0.53 kgm^2^, and *H* = 0.67 m (*M* is comparable to the measured data in ***Hudson et al***. (***2012***)), which resulted in *j* = 0.063. From the measured data of horses, we determined *M* = 245 kg, *J* = 44 kgm^2^, and *H* = 1.2 m, which resulted in *j* = 0.14. Figure 5 shows the relation of *Ψ*_1_ and *c*_1_ that satisfy Γ(*Ψ*_1_, *c*_1_) = 0 and produce the solution for various values of *d* and *k* by using the value of *j* obtained from the measured data. Specifically, we used 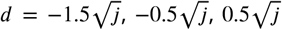, and 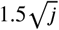 for both models, *k* = 0.5, 0.75, and 1.0 for the cheetah model, and *k* = 0.75, 1.0, and 1.25 for the horse model. The solution type depended little on *k*, and largely depended on. For 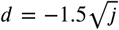 and 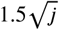, only type C and only type E, respectively, exists and the solution is unique for *Ψ*_1_ and *c*_1_. In contrast, for 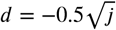 and 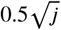, types CC and CE as well as type C and types EE and EC as well as type E exist, respectively, and the solution is not necessarily unique for *Ψ*_1_ or *c*_1_.

**Figure 5.**
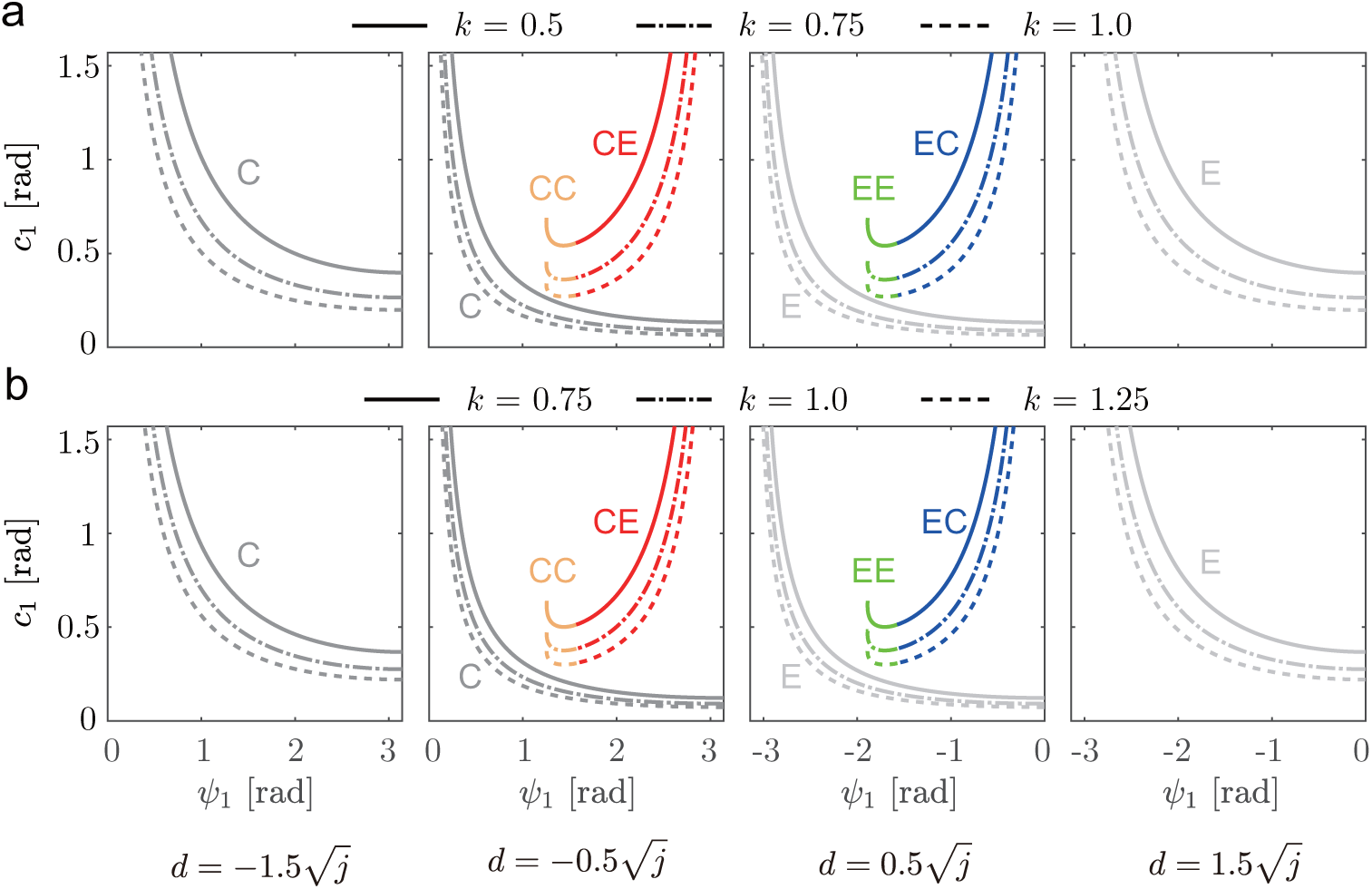
Relation of *Ψ*_1_ and *c*_1_ for periodic solutions for various *k* and *d* for (a) cheetah and (b) horse models. The solution type depends little on *k*, but largely depends on *d*.

To more clearly show the dependence of the solution type on *d*, Fig. 6a and b show the obtained solution types for *d* and *c*_1_ by projecting the solutions in the *d*-*c*_1_-*Ψ*_1_ space to the *d*-*c*_1_ plane, where we used *k* = 0.80 (*K* = 98 Nm/rad) for the cheetah model and *k* = 1.0 (*K* = 2.9× 10^3^ Nm/rad) for the horse model (*k* of the cheetah model is smaller than that of the horse model, consistent with the suggestion that cheetahs have more flexible bodies than horses in ***Hildebrand*** (***1959***)). Because the spine is never bent to the right angle during galloping, we showed the range 0 ≤ *c*_1_ ≤ *π*/2. Solutions with two identical flights appear when *d* ≠ 0. Specifically, types C and E exist for *d* < 0 and *d* > 0, respectively. In contrast, solutio<u>n</u>s with two different flights appear when 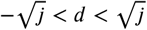. Specifically, types CC and <u>C</u>E exist for 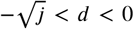, types CE and EC exist for *d* = 0, and types EE and EC exist for 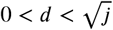.

**Figure 6.**
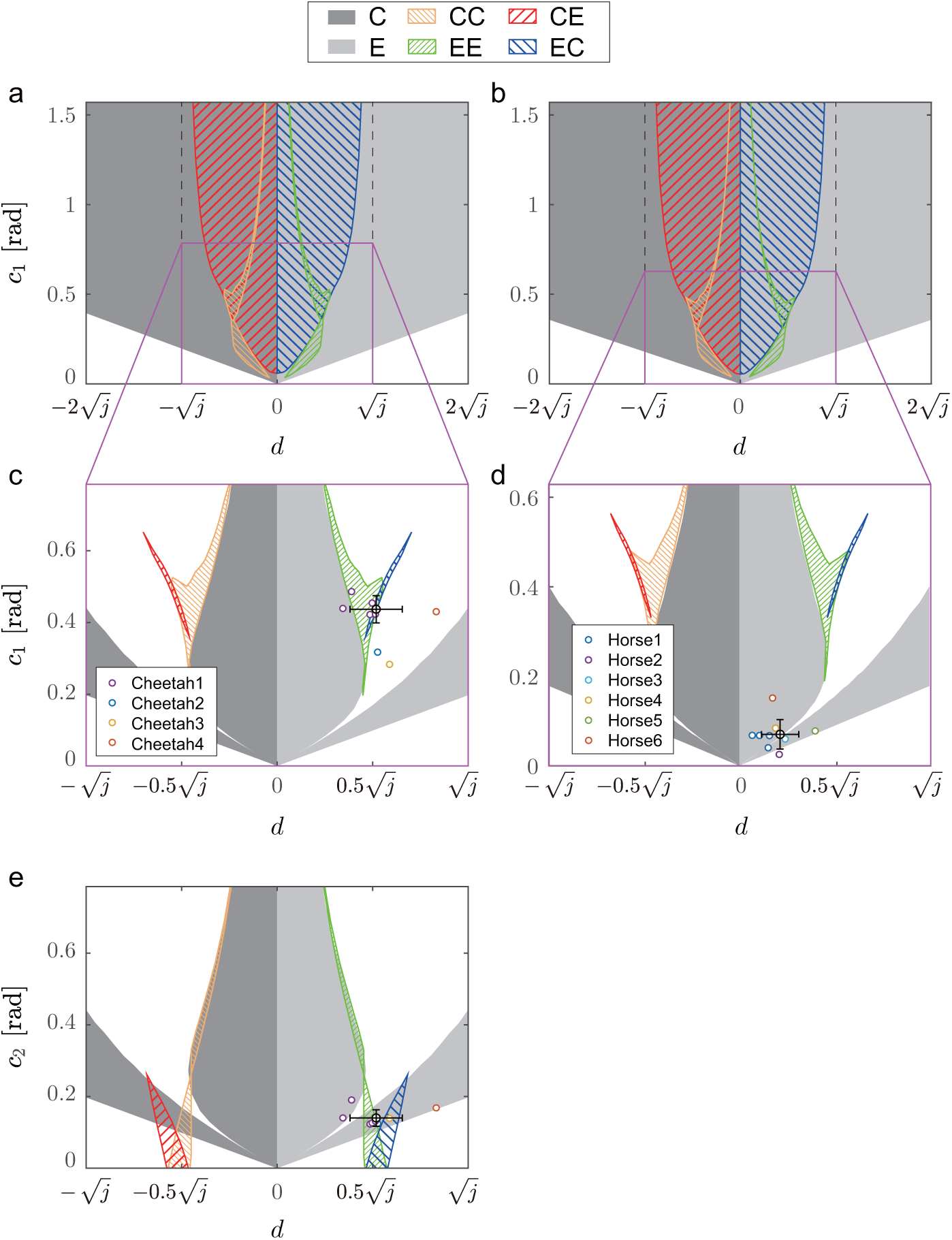
Solution type for *d* and *c*_1_ for (a) cheetah and (b) horse models. Stable solutions among solutions in a focused range of *d* and *c*_1_ in (a) and (b) for (c) cheetah and (d) horse models. (e) Stable solutions for *d* and *c*_2_ for cheetah model. Colored circles show measured animal data. Black circles and error bars show average values and standard deviations of measured animal data, respectively.

To further clarify important characteristics of the solutions from a dynamic viewpoint, Fig. 6c and d show stable solutions in a focused range of *d* and *c*_1_ among the solutions in Fig. 6a and b, respectively. Figure 6e also shows stable solutions for *d* and *c*_2_ for the cheetah model. These stable solutions were compared with the measured animal data. While type EC had a small region for stable solution, the measured data for cheetahs were located close to the stable region, as shown in Fig. 6c and e. In contrast, the measured data for horses were located in the region of stable solution of type E, as shown in Fig. 6d. Furthermore, stable solutions of type EC did not exist in the range of *d* measured from horses.

### Evaluation of the effect of GRF on spine movement

We compared the ratio of the angular velocity change *ϵ* between cheetahs and horses. The periodic solutions showed 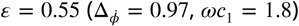 for the cheetah model and 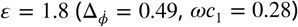 for the horse model. The measured data showed 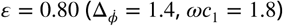 for cheetahs and 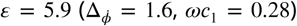 for horses. For both the periodic solutions and measured data, while *ϵ* < 1 for cheetahs, *ϵ* > 1 for horses.

### Evaluation of flight duration

The COM height changes in the first flight *δh*_1_ and second flight *δh*_2_ were obtained as

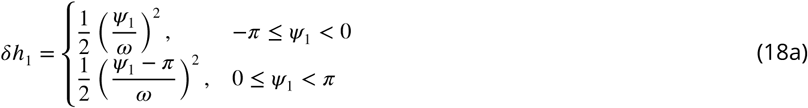

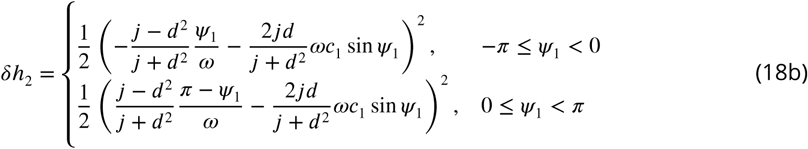

Because *δh*_1_ and *δh*_2_ monotonically decrease as *Ψ*_1_ increases when −*π* ≤ *Ψ*_1_ < 0, the COM height change of type EC is smaller than that of type E. When we compared the COM height changes between cheetahs and horses, the periodic solutions showed *δh*_1_ = 0.029 and *δh*_2_ = 0.10 for the cheetah model and *δh*_1_ = *δh*_2_ = 0.14 for the horse model. The measured data showed *δh*_1_ = 0.008 and *δh*_2_ = 0.024 for cheetahs and *δh*_1_ = *δh*_2_ = 0.12 for horses.

Changes *δΨ*_1_ and *δΨ*_2_ of the phase angle of the spine joint angle *ϕ* during the first and second flight are determined by *Ψ*_1_ and *Ψ*_2_, respectively. Specifically, for solutions of type EC, 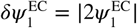 and 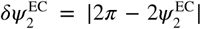, as shown in Fig. 7a, where *^*i*^ indicates the constant in the solution of type *i*. In contrast, for solutions of type E, 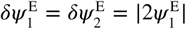, as shown in Fig. 7b. Figure 7c shows *δΨ*_1_, *δΨ*_2_, and *δΨ*_1_ + *δΨ*_2_ for the amplitude *c*_1_ of the spine joint angle *ϕ* for the cheetah model. While 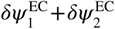 is almost 2*π* for the solution of type EC, 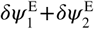 is much larger than 2*π* for the solution of type E in the range where measured data of *c*_1_ were obtained from cheetahs. The flight phase duration is determined by (*δΨ*_1_ + *δΨ*_2_)/*ω* from (16) and Fig. 4a. When *c*_1_ is identical for solutions of type EC and type E, solutions of type EC have shorter flight durations than those of type E.

**Figure 7.**
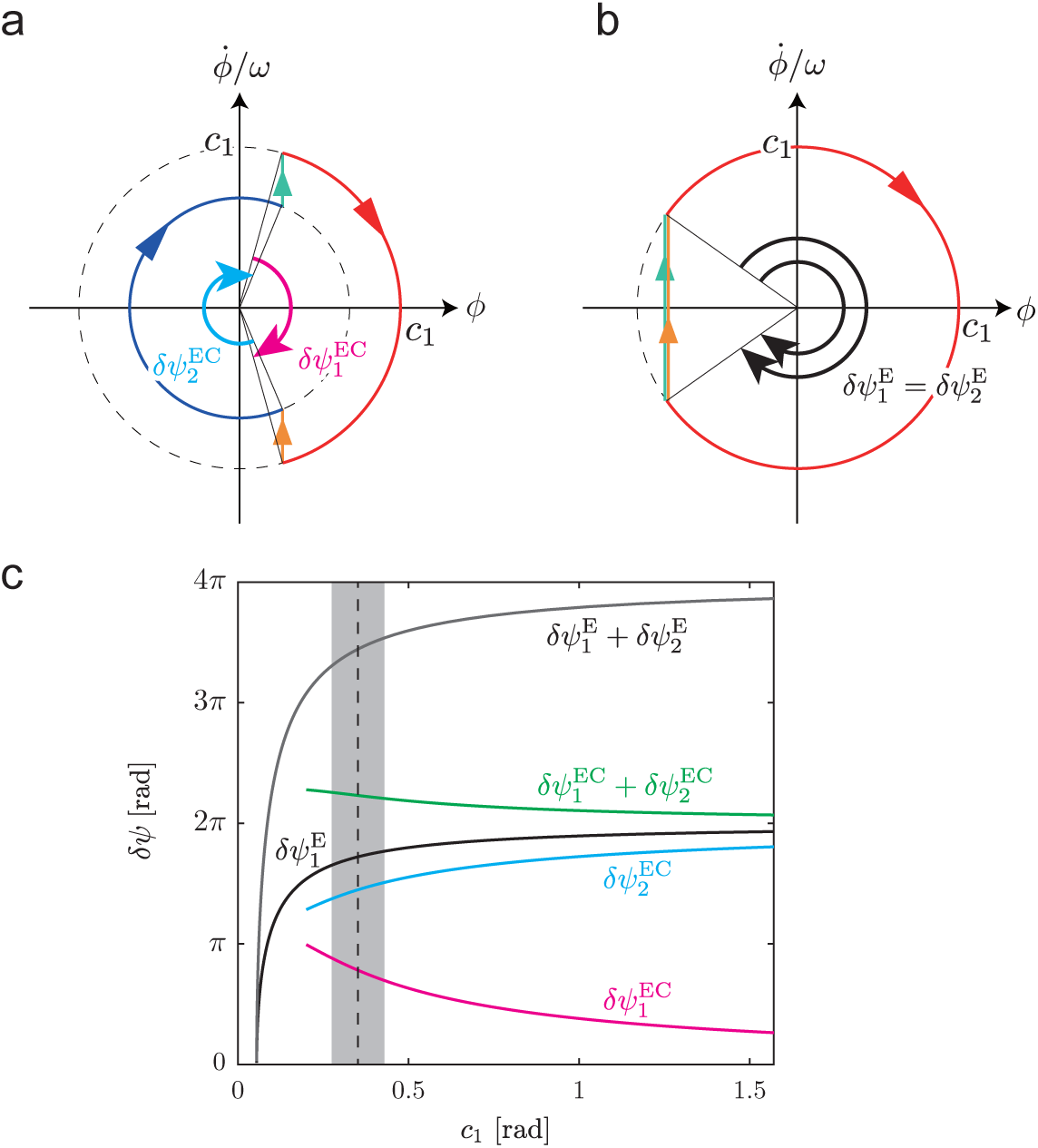
Changes *δΨ*_*i*_ (*i* = 1, 2) of phase angle of spine joint angle *ϕ* for solutions of (a) type EC and (b) type E. (c) *δΨ*_1_, *δΨ*_2_, and *δΨ*_1_ + *δΨ*_2_ for amplitude *c*_1_ of spine joint angle *ϕ* for cheetah model. Dashed line is the average of measured *c*_1_ of cheetahs, and gray area is standard deviation.

## Discussion

In the current study, to clarify the mechanisms underlying the different spine movements and gait speeds between different gallop types by focusing on the rotary gallop by cheetahs and the transverse gallop by horses from a dynamic viewpoint, we developed a simple analytical model and derived periodic solutions. The results revealed six types (C, E, CC, EE, CE, and EC) of periodic solutions (Fig. 4). These solutions were classified into two types according to their flights. While types C, E, CC, and EE involved one type of flight, types CE and EC involved two different flight types. Characteristic properties of the measured data of cheetahs were close to those of stable solutions of type EC (Fig. 6c and e). The properties of the measured data of horses were included in those of stable solutions of type E (Fig. 6d). Furthermore, there were not stable solutions of type EC in the range of *d* measured from horses (Fig. 6d).

### Mechanisms underlying different flights in rotary and transverse gallops

We investigated the different flights between the rotary and transverse gallops by focusing on cheetahs and horses. Specifically, while the rotary gallop involves two different types of flights (collected and extended), the transverse gallop involves only one type of flight (collected). Periodic solutions of types EC and CE had two different types of flights, similar to the rotary gallop. In contrast, periodic solutions of types E, C, EE, and CC involved only one type of flight, similar to the transverse gallop. We verified the obtained solutions from the comparison of the obtained solutions with measured data of animals. In particular, the impulse position *d* and the spine movement amplitude *c*_1_ and *c*_2_ of the measured data of cheetahs were close to those of the stable solutions of the type EC (Fig. 6c and e) and *d* and *c*_1_ of the measured data of horses were included in those of stable solutions of type E (Fig. 6d). Our solutions reproduced animal galloping from the view-point of spine movement. Furthermore, stable solutions of type EC did not exist in the range of *d* measured from horses (Fig. 6d), as two types of flight never appear in the transverse gallop of horses. These results suggest that the solutions of types EC and E correspond to the rotary and transverse gallop, respectively.

As described above (13), the periodic solution gave an important criterion to determine the solution type: while the signs of 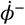 and 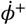 are identical for two different types of flights (including solutions of type EC), they are different for one type of flight (including solutions of type E). The criterion gave a hypothesis: while the effect of GRF is too small to change the direction of the spine movement in the cheetah rotary gallop, the effect is so large that the direction changes in the horse transverse gallop. To evaluate this hypothesis, we calculated the angular velocity change 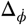 of the spine movement caused by the GRF and the amplitude *ωc*_1_ of the angular velocity, and obtained the ratio 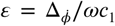 in the obtained solutions and animals. When the direction of the spine movement does not change, *ε* < 1 because 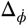 is smaller than *ωc*_1_. In contrast, when *ε* > 1, the direction changes because 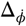 is larger than *ωc*_1_. We achieved *ε* < 1 in cheetahs while *ε* > 1 in horses for both the solutions and measured data. Furthermore, 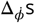 were not so different between cheetahs and horses, and *ωc*_1_ of cheetahs was much larger (over six times) than that of horses, as shown in Fig. 8. These results suggest that cheetahs have so large spine movement as to reduce the effect of the GRF, which prevents the direction from changing. This allows cheetahs to create two different flight types. In contrast, horses do not exhibit substantial spine movement, which lets the GRF change the direction. This forces horses to create only one flight type.

**Figure 8.**
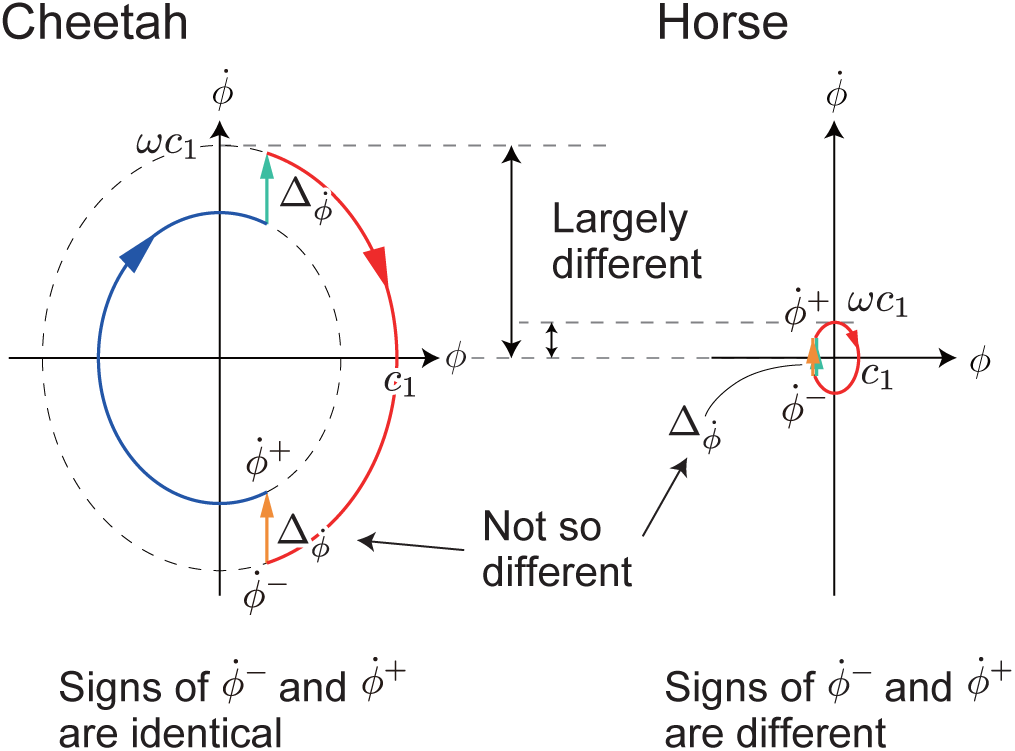
Schematics to explain the mechanism under which while cheetahs involve two different types of flights, horses involve one type of flight. 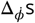 s are not so different between cheetahs and horses, and *ωc*_1_ of cheetahs is much larger than that of horses. Therefore, while signs of 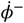 and 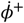 are identical for cheetahs, they are different for horses.

### Rotary gallop enables high-speed locomotion

Small gait cycle durations allow animals to kick the ground frequently for acceleration and achieve high-speed locomotion (***Hudson et al., 2012***). As shown below (18), COM height changes *δh*_1_ and *δh*_2_ of cheetahs are both smaller than those of horses, which implies that the flight durations of cheetahs are smaller than those of horses. This result suggests that cheetahs use rotary gallop to create smaller flight durations than those of the horse transverse gallop, which allows cheetahs to move faster than horses.

The criterion explained above also provided the mechanism by which the rotary gallop produces higher-speed locomotion than the transverse gallop. When cheetahs show both solutions of type EC and type E with identical *c*_1_, solutions of type EC have shorter flight phases than those of type E, as shown in Fig. 7c. This is because while the phase angle change *δΨ*_1_ + *δΨ*_2_ of the solutions of type EC is almost 2*π*, that of the solutions of type E is much larger than 2*π*, which makes the gait cycles of the solutions of type EC shorter than those of the solutions of type E. This result suggests that cheetahs produce high-speed locomotion using rotary gallop rather than transverse gallop.

### Parameter dependence of solutions

The type of periodic solutions depended on the relationship between *d* and *j* (Figs. 5 and 6). In particular, the solutions of types CE and EC existed only when 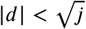. The foot-contact relationship (6) plays an important role in this parameter dependence. We explain this mechanism below. More detailed explanations are provided in Supplementary information S4.

#### Why solutions of types EC and CE exist when 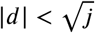

First, we suppose that 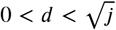. When the first flight is extended, −*π* ≤ *Ψ*_1_ < 0 is obtained from (7) because 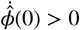. The substitution of (7) into the fourth row of (10) gives

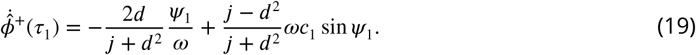

Because the first and second terms of the right-hand side are positive and negative, respectively, the sign of 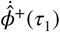 and the type of the second flight depend on *Ψ* and *c*. When 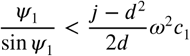 or when 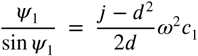 and 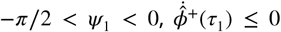 is satisfied and the second flight is collected. When 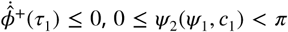 is obtained from (7). The substitution of (7) into (17) gives

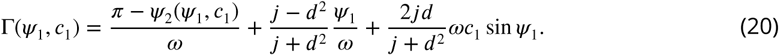

In contrast, (7) gives

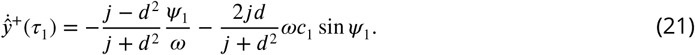

Because 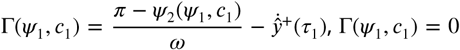 can be satisfied when 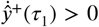. Therefore, periodic solutions of type CE can exist. When 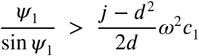 or when 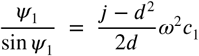 and 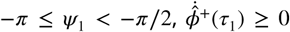 is satisfied and the second flight is extended. When 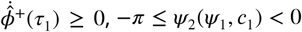 is obtained from (7). The substitution of (7) into (17) gives

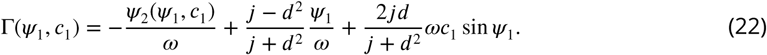

Because 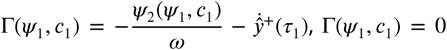 can be satisfied when 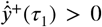. Therefore, periodic solutions of types E and EE can exist.

Second, we suppose that 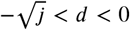. In this case, we can show that the solutions of type CE can exist, in the same way as above (because the second flight can be both extended and collected when the first flight is collected, solutions of types C and CC can also exist).

#### Why solutions of types EC and CE never exist when 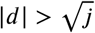

Next, we suppose that 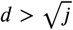. When the first flight is extended, 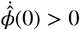 is satisfied and −*π* ≤ *Ψ*_1_ < 0 is obtained from (7). The substitution of (7) and (16e) into the fourth row of (10) gives

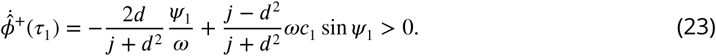

This indicates that the second flight is also extended and that solutions of type EC never exist.

Finally, we suppose that 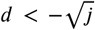. In this case, we ca<u>n</u> show that solutions of type CE never exist, in the same way as above. Therefore, when 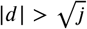, the solutions of type EC and CE never exist due to the foot-contact relationship.

### Limitations and future work

In the current study, the results revealed that our simple model has six types of solutions and that the change of angular velocity of the spine joint by the vertical impulse determines the type of solution obtained. The obtained periodic solutions had similar characteristics to the measured data of cheetahs and horses. In particular, stable solutions of type EC reproduced the rotary gallop of cheetahs from the viewpoint of spine movement. However, our model was limited in its ability to accurately reproduce galloping of horses. For example, although the transverse gallop of horses involves only collected flight, the measured data were located in the stable solutions of type E but not type C (Fig. 6d). To overcome this limitation, several improvements are needed. For example, our two rigid bodies need to be asymmetric because horses have different physical properties between the fore and hind parts of the body, and bend their backs around the sacrum rather than in the middle of the spine (***Hildebrand, 1959***). In addition, the torsional spring in the spine joint should be asymmetric in the extension and flexion directions because the spine of the horse is difficult to bend (***Gyambaryan, 1974***), particularly in the extending direction (***Licka and Peham, 1998***). Furthermore, we neglected the dynamics in the horizontal and pitching direction in our model. It has also been suggested that cheetahs achieve high-speed locomotion by extended flight, so that the touchdown of the forelimbs does not decelerate in the horizontal direction (***Bertram and Gutmann, 2009***). We would like to improve our model to investigate these effects in future research.

Our model included a torsional spring connecting two rigid bodies. Previous animal measurement data suggest that animals use their bodies as elastic structures, such as the tendons in the torso (***Taylor, 1978***; ***Alexander, 1988***). However, trunk muscles are also effectively used as actuators to produce energy for acceleration (***Hildebrand, 1959***; ***Biancardi and Minetti, 2012***). Finally, we also intend to investigate the effect of trunk control on locomotion speed and energy efficiency in the future.

## Acknowledgments

This work was supported in part by JSPS KAKENHI Grant Number JP18J10682. The authors thank all of the staff at Shanghai Wild Animal Park for their support and help during data collection. The authors thank Taiki Matsuo in The United Graduate School of Veterinary Science, Yamaguchi University, Yamaguchi, Japan, for collecting CT data of cheetah. The authors thank all of the staff at the JRA Equine Research Institute for their generous assistance with the measurement of the horse.

## Supplementary files

### Supplementary file 1

(S1) Effect of assumption on galloping dynamics. (S2) Foot contact dynamics. (S3) Role of symmetry condition in solution. (S4) Parameter dependence of solutions.

**Supplementary file 1**

(S1) Effect of assumption on galloping dynamics. (S2) Foot contact dynamics. (S3) Role of symmetry condition in solution. (S4) Parameter dependence of solutions.

## S1 Effect of assumption on galloping dynamics

We ignored the pitching movement of the whole body in our model (Fig. 2) because the COM vertical and spine joint movements are more important for determining galloping dynamics, compared with pitching movements. This assumption induced simultaneous foot contact between the fore and hind legs. We investigated this dynamical effect based on a model which incorporates *θ* as the pitch angle of the whole body, as shown in Fig. S1. In this case, the foot contact does not necessarily occur simultaneously between the fore and hind legs.

The motion of this model is governed by the equations of motion of *Y, θ*, and *ϕ*, which are given by

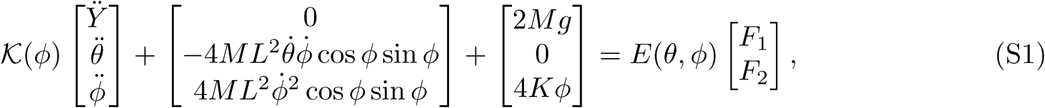

where

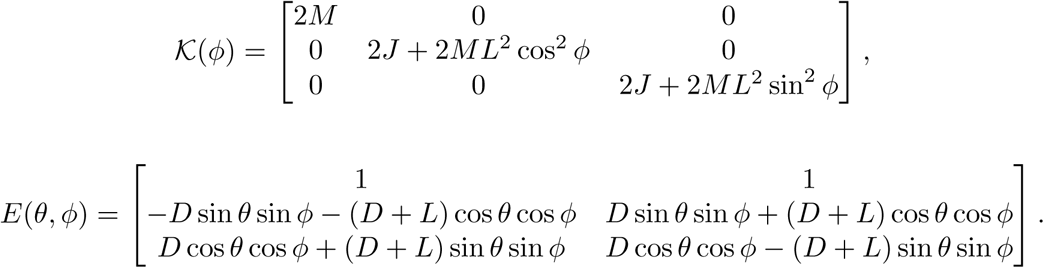

*F*_1_ and *F*_2_ are the vertical reaction forces of the fore and hind legs, respectively (*F*_*i*_ > 0 for the stance phase, *F*_*i*_ = 0 for the swing phase). The relationship between the states immediately prior to and immediately following the foot contact is given by

**Fig S1.**
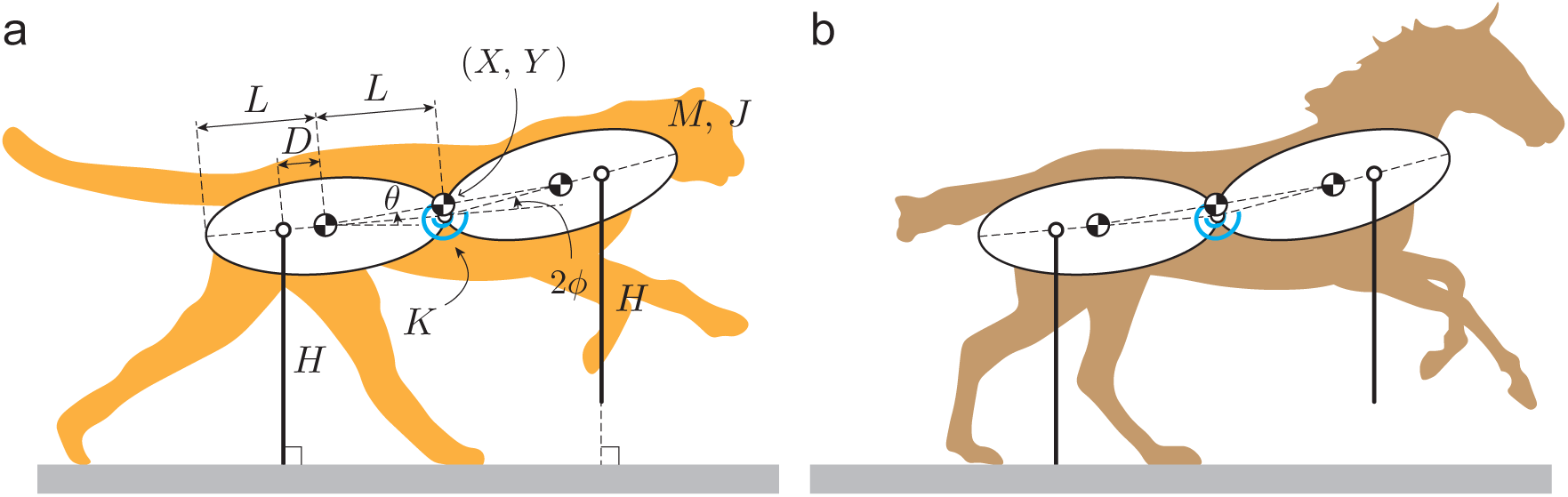
Models for (a) cheetah and (b) horse, which incorporate the pitch angle *θ* of the whole body. Foot contacts do not necessarily occur simultaneously between the fore and hind legs. Fig. S1 Models for (a) cheetah and (b) horse, which incorporate the pitch angle *θ* of the whole body. Foot contacts do not necessarily occur simultaneously between the fore and hind legs.

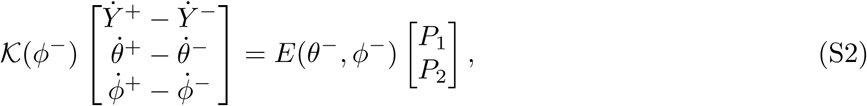

where *P*_1_ and *P*_2_ are the impulses of the fore and hind legs, respectively (*P*_1_ > 0 for the foot contact of the fore leg, *P*_2_ > 0 for the foot contact of the hind leg; otherwise *P*_*i*_ = 0), and can be determined to satisfy the energy conservation.

We assumed that 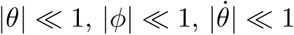,. and 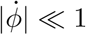. The linearization of the equations of motion (S1) and relationship between the states immediately prior to and immediately following the foot contact (S2) gives

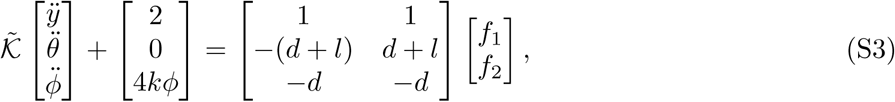

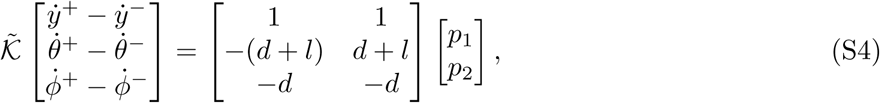

where

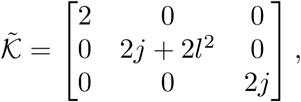

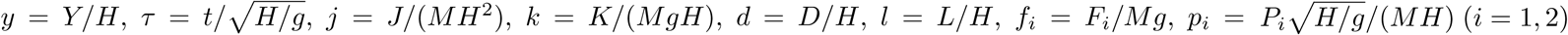, and 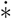 indicates the derivative of variable ∗ with respect to *τ*. These equations are reduced to each equation of *y, θ*, and *ϕ*. While *f*_1_ and *f*_2_ (*p*_1_ and *p*_2_) have different effects (opposite signs) on the equation of *θ*, they have the same effect on each equation of *y* and *ϕ* (same sign). Therefore, the following three cases *f*_1_ = *f*_2_ = *f* (*p*_1_ = *p*_2_ = *p*), *f*_1_ = 2*f* and *f*_2_ = 0 (*p*_1_ = 2*p* and *p*_2_ = 0), and *f*_1_ = 0 and *f*_2_ = 2*f* (*p*_1_ = 0 and *p*_2_ = 2*p*) have the same dynamic effect on *y* and *ϕ*.

When *θ* = 0, the motions of the fore and hind parts of the model are symmetrical, resulting in simultaneous foot contact between the fore and hind legs and *f*_1_ = *f*_2_ = *f* (*p*_1_ = *p*_2_ = *p*). This effect on *y* and *ϕ* is identical to that of individual foot contact between fore and hind legs with *f*_1_ = 2*f* and *f*_2_ = 0 (*p*_1_ = 2*p* and *p*_2_ = 0) and *f*_1_ = 0 and *f*_2_ = 2*f* (*p*_1_ = 0 and *p*_2_ = 2*p*). Therefore, even when we ignore the pitching movement (*θ* = 0), *y* and *ϕ* have no significant effect.

## S2 Foot contact dynamics

Here, we derive the relationship (3) between the states immediately prior to and immediately following foot contact in the model. We assumed elastic collision for foot contact that involves no position change and energy conservation. We define Δ_*P*_ as the impulse at foot contact from the ground in the vertical direction. 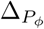 is the change in the angular momentum caused by the impulse. The relationship of the translational and angular momentum between immediately prior to and following the foot contact gives

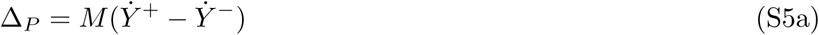

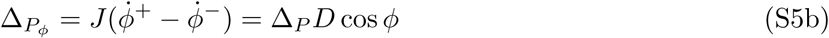

From energy conservation, we obtain

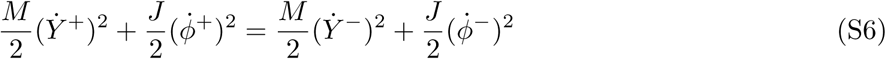

From (S5) and (S6), we obtain (3).

## S3 Role of symmetry condition in solution

Here, we show the mechanism by which the symmetry condition (12) forces the third and fourth rows in (11) to be satisfied. The substitution of (12) into the first row of (10) gives

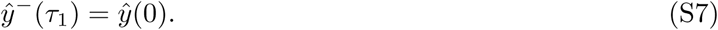

By substituting (7a) into (S7), we obtain

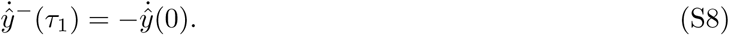

By substituting the first and second rows of (11) into (9), we obtain

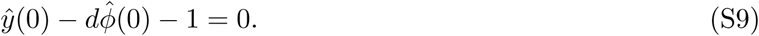

In contrast, by substituting (S7) into (8), we obtain

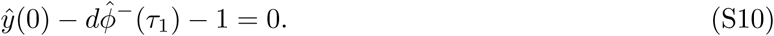

(S9) and (S10) give

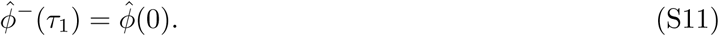

Because we assumed *τ*_1_ < 2*π/ω*, the substitution of (7b) into (S11) gives

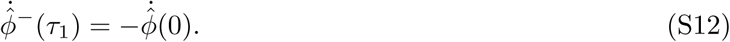

Therefore, from (S7), (S8), (S11), and (S12), the relationship between the states at the beginning of the first flight phase (*τ* = 0) and immediately prior to the first foot contact (*τ* = *τ*_1_) is given by

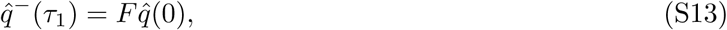

where

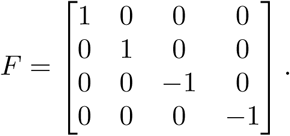

By substituting (S11) into the second row of (10), we obtain

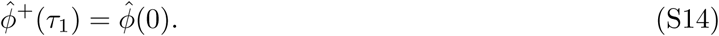

The first and second rows of (11), (12), and (S14) give

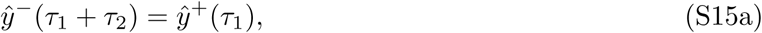

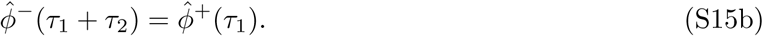

Because we assume *τ*_2_ < 2*π/ω*, the substitution of (7) into (S15) gives

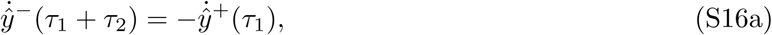

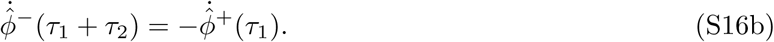

Therefore, from (S15) and (S16), the relationship between the states immediately following to the first foot contact (*τ* = *τ*_1_) and immediately prior to the second foot contact (*τ* = *τ*_1_ + *τ*_2_) is also given by

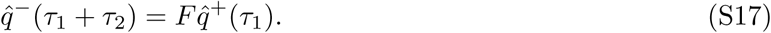

From (10), (S13), and (S17), we obtain

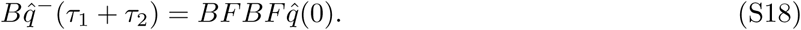

Because *BFBF* = *I*, where *I* is an identity matrix, 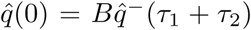 is satisfied. This is equal to (11). Therefore, the third and fourth rows of (11) are satisfied when the symmetry condition (12) is given.

## S4 Parameter dependence of solutions

Here, we show how the types of solutions depend on *d* and *j*.

### S4.1 When 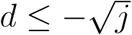

When the first flight is collected, 0 ≤ *ψ*_1_<*π* is obtained from (7) because 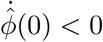. The substitution of (7) into the fourth row of (10) gives

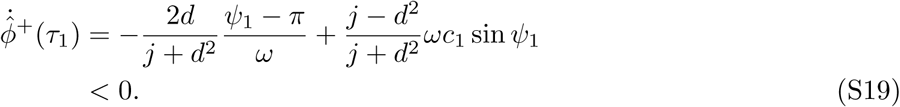

This indicates that the second flight is also collected. Therefore, periodic solutions of type CE never exist. From (7) and (S19), 0 ≤ *ψ*_2_(*ψ*_1_, *c*_1_) <*π* is obtained and (16) gives

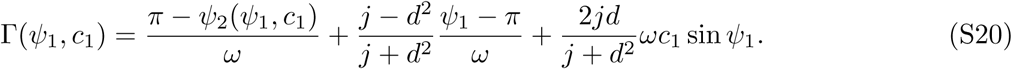

In contrast, (7) gives

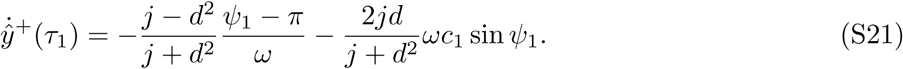

Because 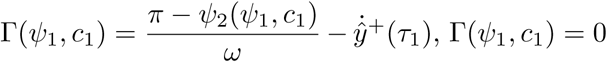 can be satisfied when 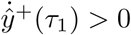. Therefore, periodic solutions of types C and CC can exist.

When the first flight is extended, −*π* ≤ *ψ*_1_< 0 is obtained from (7) because 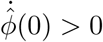. The substitution of (7) into the fourth row of (10) gives

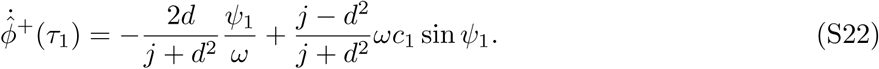

Because the first and second terms of the right-hand side are positive and negative, respectively, the sign of 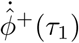 and the type of the second flight depend on *ψ*_1_ and *c*_1_. When 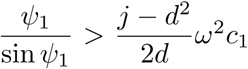 or when 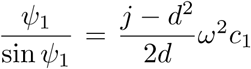 and 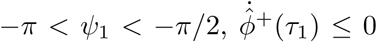 is satisfied and the second flight is collected. However, 0 ≤ *ψ*_2_(*ψ*_1_, *c*_1_) <*π* is obtained from (7). The substitution of (7) into (16) gives

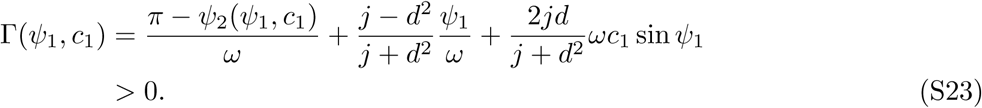

Because Γ(*ψ*_1_, *c*_1_) = 0 is not satisfied, solutions of type EC never exist. When 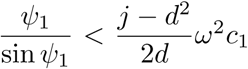 or when –*π/*2 < *ψ*_1_< 0 and 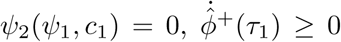 is satisfied and the second flight is extended. However, −*π* ≤ *ψ*_2_(*ψ*_1_, *c*_1_) < 0 is obtained from (7). The substitution of (7) into (16) gives

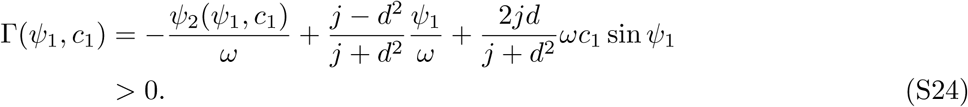

Because Γ(*ψ*_1_, *c*_1_) = 0 is not satisfied, solutions of types E and EE never exist. When 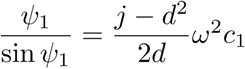 and 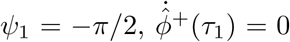 is satisfied. In this case, *c*_2_(*ψ*_1_, *c*_1_) =0 is obtained from (14d). Therefore, the second flight is neither extended nor collected (Fig. S2a). However, the substitution of 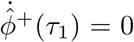 and (7) into (16) gives

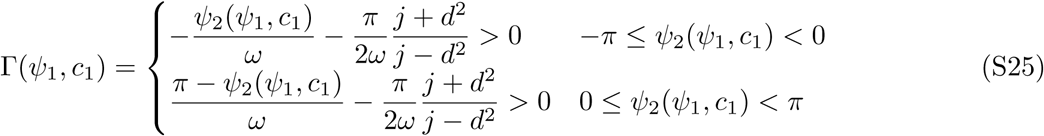

Because Γ(*ψ*_1_, *c*_1_) = 0 is not satisfied, solutions like Fig. S2a never exist.

Therefore, only the solutions of types C and CC can exist when 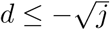.

**Fig. S2.**
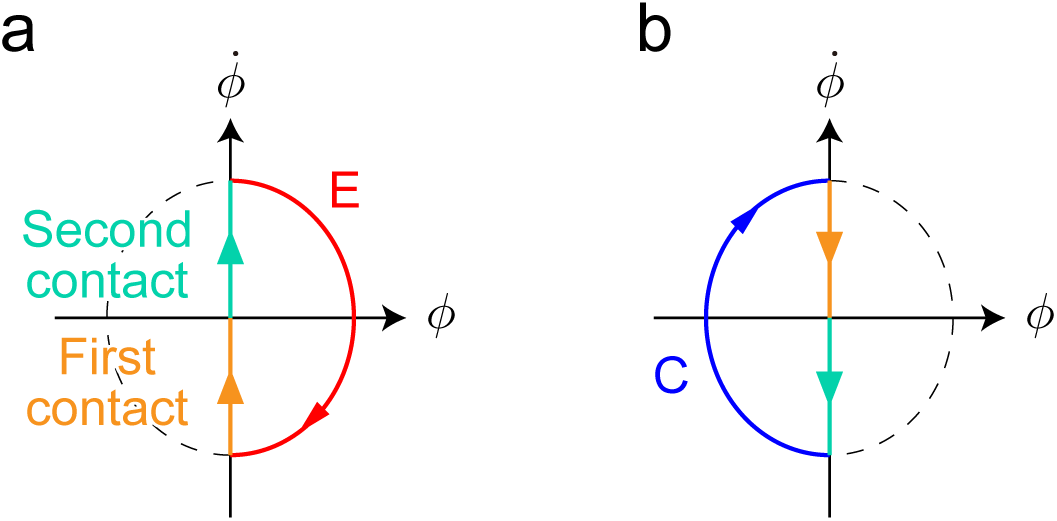
Periodic solutions whose second flight is neither extended nor collected. (a) *ψ*_1_ = *−π/*2. (b) *ψ*_1_ = *π/*2.

### S4.2 When 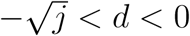

When the first flight is collected, 0 ≤ *ψ*_1_< *π* is obtained from (7) because 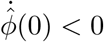. The substitution of (7) into the fourth row of (10) gives

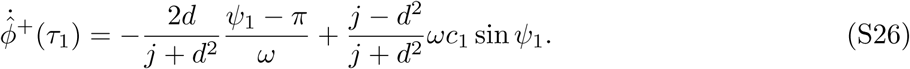

Because the first and second terms of the right-hand side are negative and positive, respectively, the sign of 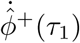 and the type of the second flight depends on *ψ*_1_ and *c*_1_. When 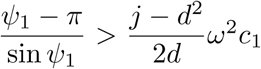 or when 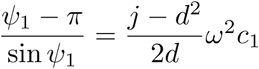 and 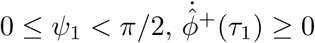 is satisfied and the second flight is extended. − *π* ≤ *ψ*_2_ (*ψ*_1_, *c*_1_) < 0 is obtained from (7) because 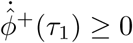. The substitution of (7) into (16) gives

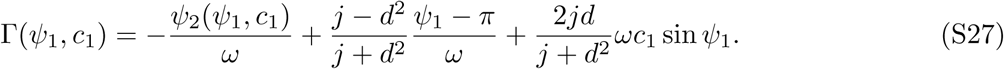

In contrast, (7) gives

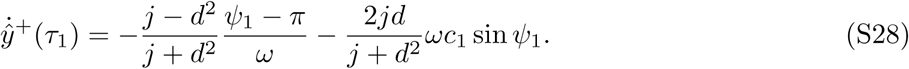

Because 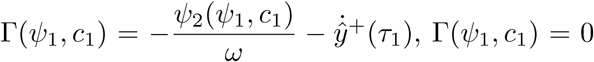 can be satisfied when 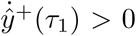. Therefore, periodic solutions of type CE can exist. When 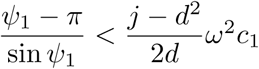 or when 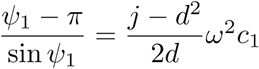 and 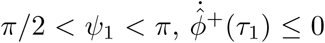 is satisfied and the second flight is collected. 0 ≤ *ψ*_2_(*ψ*_1_, *c*_1_) < *π* is obtained from (7) because 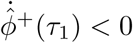. The substitution of (7) into (16) gives

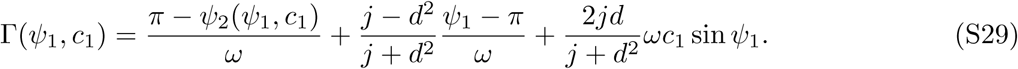

In contrast, (7) gives

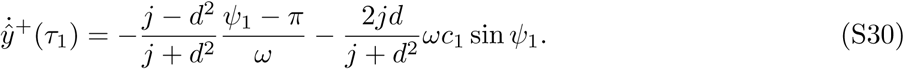

Because 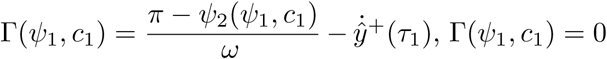 can be satisfied when 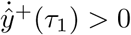. Therefore, periodic solutions of types C and CC can exist. When 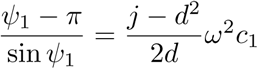 and 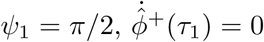 is satisfied. In this case, *c*_2_(*ψ*_1_, *c*_1_) = 0 is obtained from (7). Therefore, the second flight is neither extended nor collected (Fig. S2b). However, the substitution of 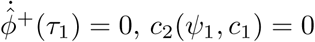, and (7) into (16) gives

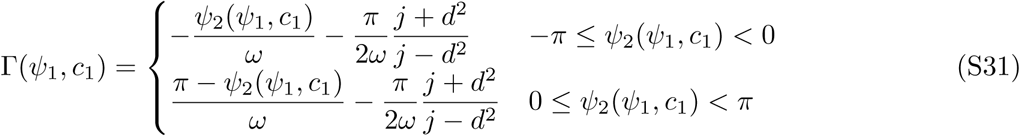

Γ(*ψ*_1_, *c*_1_) = 0 is satisfied only when 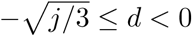. Therefore, periodic solutions like Fig. 2b exist only when 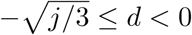.

When the first flight is extended, −*π* ≤ *ψ*_1_< 0 is obtained from (7) because 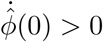. The substitution of (7) into the fourth row of (10) gives

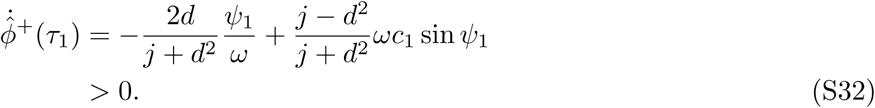

This indicates that the second flight is collected. Therefore, periodic solutions of types E and EE never exist. From (7) and (S32), 0 ≤ *ψ*_2_(*ψ*_1_, *c*_1_) <*π* is obtained. The substitution of (7) into (16) gives

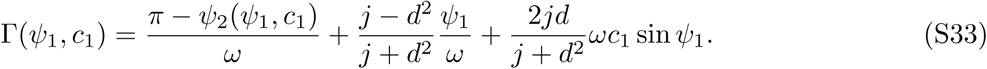

In contrast, (7) gives

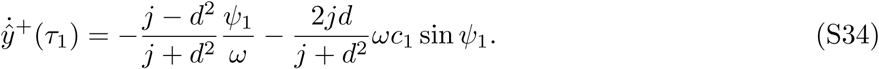

Because 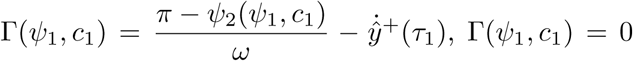 can be satisfied when 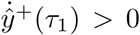. From 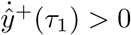, we obtain

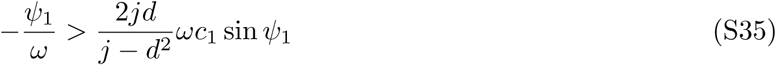

From the substitution of (S35) into (15d), we obtain

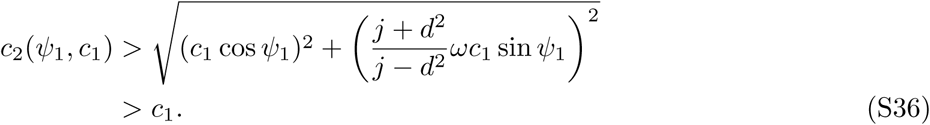

Because we assumed *c*_1_ > *c*_2_ in Section 2.3, solutions of type EC never exist.

Therefore, only the solutions of types C, CC, and CE, and solutions like Fig. S2b exist when 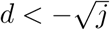.

### S4.3 When *d* =0

From (7), *a*_2_(*ψ*_1_, *c*_1_) = *a*_1_(*ψ*_1_), *c*_2_(*ψ*_1_, *c*_1_) = *c*_1_, and *ψ*_2_(*ψ*_1_, *c*_1_) = *ψ*_1_ are obtained. When the first flight is extended, −*π* ≤ *ψ*_1_< 0 is obtained from (7) because 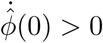. The substitution of (7) into the fourth row of (10) gives

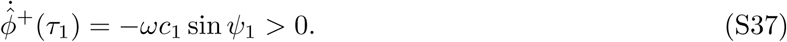

This indicates that the second flight is collected. Therefore, solutions of types E and EE never exist. The substitution of (7) into (16) gives

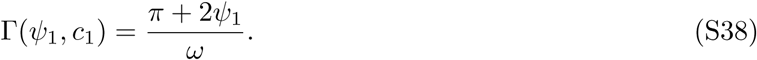

Γ(*ψ*_1_, *c*_1_) = 0 is satisfied when *ψ*_1_ = −*π/*2. Therefore, solutions of type EC exist.

When the first flight is collected, 0 ≤ *ψ*_1_<*π* is obtained from (7) because 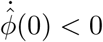. The substitution of (7) into the fourth row of (10) gives

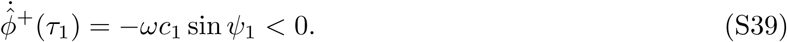

This indicates that the second flight is extended. Therefore, solutions of types C and CC never exist. The substitution of (7) into (16) gives

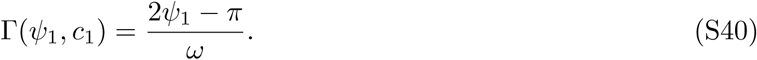

Γ(*ψ*_1_, *c*_1_) = 0 is satisfied when *ψ*_1_ = *π/*2. Therefore, solutions of type CE exist.

Therefore, only solutions of types CE or EC exist (they are identical because *c*_1_ = *c*_2_).

### S4.4 When 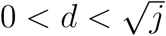

Only solutions of types E, EE, and EC can exist in the same way as S4.2.

### S4.5 When 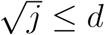

Only solutions of types E and EE can exist in the same way as S4.1.

